# Structural Compression and Entorhinal Vulnerability: Linking Tentorial Adjacency to Tau Burden and Dementia Progression

**DOI:** 10.1101/2025.09.02.670249

**Authors:** Luyue Zhang, Ana M. Franceschi, John F. Crary, Frank A. Provenzano, the Alzheimer’s Disease Neuroimaging Initiative

**Affiliations:** Department of Biomedical Engineering, Columbia University New York, NY, USA; Neuroradiology Division, Department of Radiology, Donald and Barbara Zucker School of Medicine at Hofstra/Northwell, Lenox Hill Hospital, New York, NY, USA; Department of Pathology, Nash Family Department of Neuroscience, Department of Artificial Intelligence & Human Health, Neuropathology Brain Bank & Research CoRE, Ronald M. Loeb Center for Alzheimer’s Disease, Friedman Brain Institute, Icahn School of Medicine at Mount Sinai, New York, NY, USA; Department of Neurology, Columbia University Irving Medical Center, New York, NY, USA

**Keywords:** Tau PET, aging, Alzheimer’s disease, neuropathology, entorhinal cortex, Tauopathy, mild traumatic brain injury

## Abstract

Alzheimer’s disease (AD) is a growing public health crisis. The disease is defined neuropathologically by accumulation of amyloid-β plaques and neurofibrillary tangles (NFTs) composed of abnormal tau protein in the brain. Early neurofibrillary degeneration in the entorhinal cortex (EC) is a hallmark of AD and a critical initiating event in the hierarchical pathoanatomical progression. However, the factors triggering initial tau deposition in the EC remain unclear. We propose a novel biomechanical cascade hypothesis, positing that the unique anatomical inferomedial positioning of the EC, including proximity to the tentorial incisura (TI) and other skull base structures, renders it susceptible to very mild yet persistent age-related mechanical stress, analogous to the effects of repetitive mild traumatic brain injury, triggering tau pathology. To test this hypothesis, we developed a method to quantify Entorhinal-Tentorial (EC-TI) proximity and applied it to multimodal imaging data from the Alzheimer’s Disease Neuroimaging Initiative (ADNI; *n*=47). Based on this neuroanatomical contact coefficient (NCC), participants were heuristically stratified into high (*n*=24) and low (*n*=23) adjacency groups. When controlling for other risk factors, tau PET signal in the EC predicted conversion from mild cognitive impairment to AD only in the high-adjacency group (LLR *p*=0.009, tau PET in EC *p*=0.036). These findings identify EC-TI proximity as a novel and anatomically grounded biomarker of AD progression risk. More broadly, they suggest a previously unrecognized biomechanical contribution to the initiation of tau pathology in aging and sporadic AD, opening new avenues for early detection, risk stratification, and mechanistically targeted prevention strategies.

## INTRODUCTION

Deposition of neurofibrillary tangles composed of the microtubule-associated protein tau are a hallmark of a diverse group of disorders termed the tau proteinopathies (“tauopathies”), with Alzheimer’s disease (AD) being the most common (Lee et al., 2001; Soto, 2003). In early AD, tau deposition is prominent in the entorhinal cortex, a key structure within the hippocampal formation in the medial temporal lobe (Braak & Braak, 1991). Tauopathic changes are thought to progress over time, ultimately disrupting regional brain network dynamics (Kocagoncu et al., 2020).

However, tau accumulation in the temporal lobe is not exclusive to AD. Primary age-related tauopathy (PART), an amyloid-independent neurodegenerative process, also features tau pathology confined predominantly to the medial temporal lobe (Crary et al., 2014). Unlike AD, PART lacks significant amyloid-β deposition and primarily affects elderly individuals, highlighting the complexity of tauopathies beyond amyloid-driven or related pathogenic processes.

The mechanism of the early selective vulnerability of the entorhinal cortex remains elusive, suggesting activity on a cellular layer level may lead to degeneration, with no clear specific cell types appearing to drive neurodegeneration (Zhao et al., 2022). The neurons in layer II of the entorhinal cortex are believed to be the initially vulnerable EC cells (Gómez-Isla et al., 1996), and they may have specific developmental, morphological and biochemical features that contribute to their vulnerability. Vascular factors, such as hypoperfusion have also been proposed with some studies suggesting a negative correlation between blood flow and cognitive decline (Khan et al., 2014) and as well as preceding tau deposition (Kapadia et al., 2023). Another hypothesis implicates high metabolic demands, as the entorhinal region functions as a major hub for integrating sensory information; its extensive connectivity may impose metabolic stress that heightens vulnerability (Mandino et al., 2022). Inflammatory processes are also increasingly recognized as contributors to early pathology, possibly through microglial activation or cytokine-mediated signaling cascades that exacerbate neuronal dysfunction (Chen & Yu, 2023). Genetic factors may further modulate regional vulnerability through differential expression of proteins associated with neurodegeneration (Mathys et al., 2024). Lastly, the accumulation of amyloid-β in and around the entorhinal cortex may compound these processes and contribute to disease onset (Cai et al., 2023).

Tau proteinopathy is also observed as a late event following mild yet repetitive traumatic brain injury (TBI). While TBI of varying severities shows acute symptomatologies, it can lead to long-term, widespread neuropathological and cognitive complications (McKee et al., 2023). This pathological link connects TBI to an earlier onset of AD (Mendez et al., 2015), an increased dementia risk by two- to fourfold (Shively et al., 2012), and accelerated AD progression in animal models (Tran et al., 2011).

The early accumulation of tau in TBI shares mechanistic similarities with chronic traumatic encephalopathy (CTE), where repetitive mechanical trauma drives progressive tau pathology (Gavett et al., 2011). Clinically, distinguishing between CTE and AD, particularly AD cases with a history of neurotrauma, can be challenging due to overlapping symptoms and pathological features (Shively et al., 2012). However, the spatiotemporal patterns of tau deposition differ between these conditions. In CTE, tau pathology spreads to the hippocampus and entorhinal cortex during stage III, whereas in AD, these regions are affected in the earliest stages of the disease (McKee et al., 2015).

While external mechanical trauma, such as TBI, has been implicated in initiating tau propagation in AD (Hicks et al., 2022), the role of intrinsic mechanical forces in tau accumulation remains largely unexplored. One potential mechanism involves chronic mechanical stress from internal anatomical structures, leading to shearing forces and localized tau deposition. The entorhinal cortex, the initial site of tau accumulation in AD, is positioned adjacent to the rigid cerebellar tentorial wall, making it theoretically susceptible to mechanical compression (Figure 1). However, there is a marked degree of interindividual variability in the size of the tentorial opening which would be predicted to modulate the degree of compression (Adler & Milhorat, 2002).

**Figure 1.**
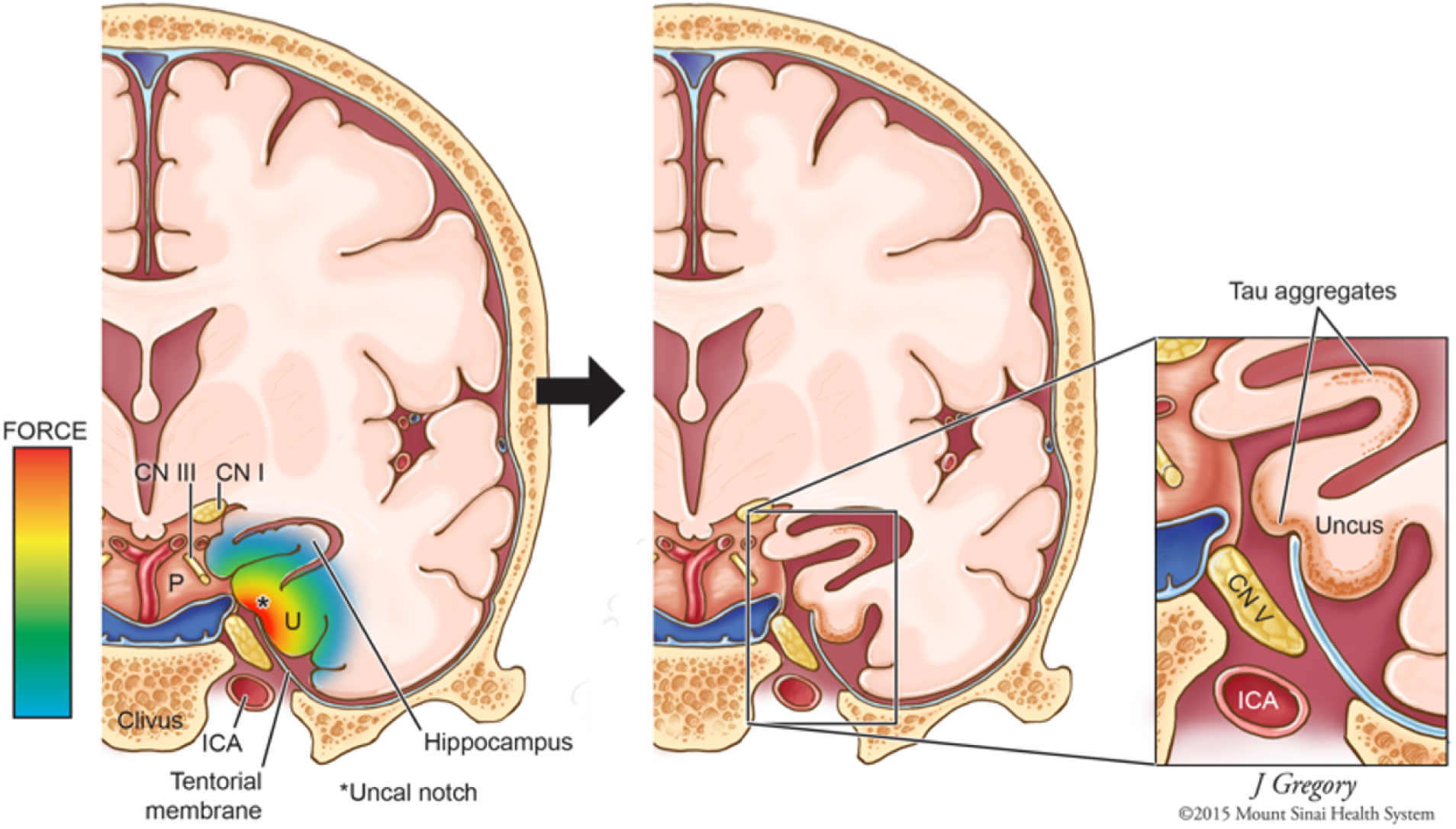
Entorhinal Cortex at Risk of Mechanical Forces by Tentorial Incisura

To investigate this hypothesis, we utilized multimodal imaging including tau PET and structural magnetic resonance imaging (MRI) in a cohort followed longitudinally. Our study sought to spatially quantify tau deposition in relation to anatomical sites of mechanical stress in the temporal lobe. Specifically, we examine how tau accumulation in EC can affect the conversion from mild cognitive impairment (MCI) to AD in patients with high and low Entorhinal-Tentorial (EC-TI) proximity defined in MRI respectively, and test whether tau accumulation correlates with impingement against the TI. These findings could offer novel insights into the potential role of chronic mechanical forces in AD pathogenesis.

## METHODS

### Subject Admission and Clinical Data Assessments

All participants were drawn from the Alzheimer’s Disease Neuroimaging Initiative (ADNI) database (<u>adni.loni.usc.edu</u>) (Alzheimer’s Disease Neuroimaging Initiative, 2004), which offers open source neuroimaging and clinical data from elderly individuals at risk for Alzheimer’s Disease. We identified and selected 47 participants with a baseline diagnosis of MCI, whose MRI and PET scans met ADNI quality control standards. Inclusion criteria required availability of T1-weighted structural MRI, tau PET scans (UC Berkeley - Tau PET 6mm Res), and amyloid PET-based status (UC Berkeley - Amyloid PET 6mm Res), all collected within 12 months of baseline MRI acquisition. Of all individuals, 24 converted to AD within 36 months of initial MCI diagnosis, while 23 remained stable. Demographic data, including age, sex, and MMSE scores, were acquired at baseline (Table 1).

**Table 1.**
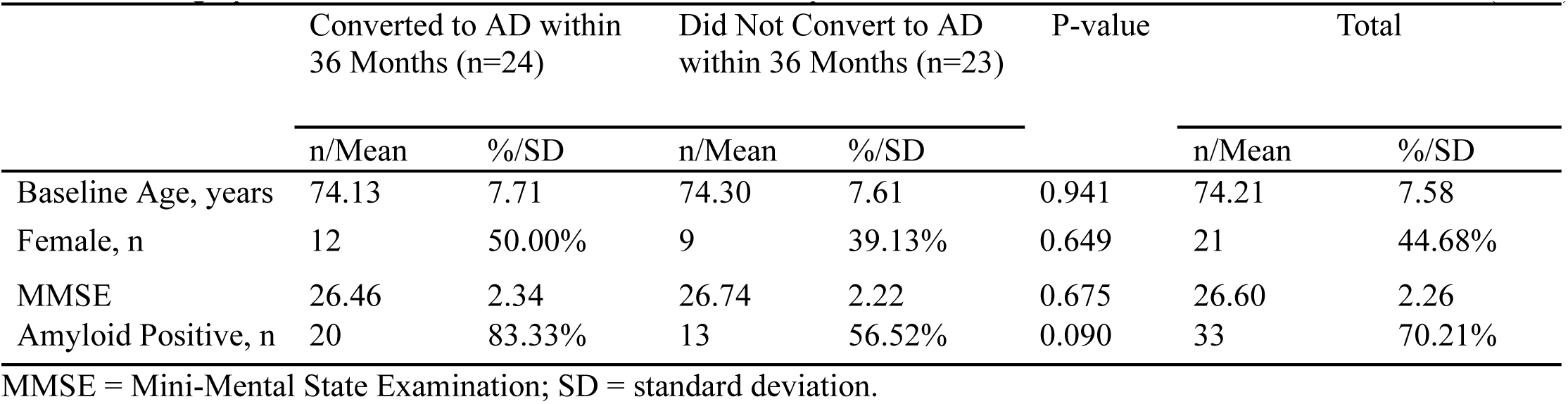
Demographic Data and Clinical Assessment of Participants that Did or Did Not Convert from MCI to AD (n=47)

|  | Converted to AD within<br>36 Months (n=24) |  | Did Not Convert to AD<br>within 36 Months (n=23) |  | P-value | Total |  |
| --- | --- | --- | --- | --- | --- | --- | --- |
|  | n/Mean | %/SD | n/Mean | %/SD |  | n/Mean | %/SD |
| Baseline Age, years | 74.13 | 7.71 | 74.30 | 7.61 | 0.941 | 74.21 | 7.58 |
| Female, n | 12 | 50.00% | 9 | 39.13% | 0.649 | 21 | 44.68% |
| MMSE | 26.46 | 2.34 | 26.74 | 2.22 | 0.675 | 26.60 | 2.26 |
| Amyloid Positive, n | 20 | 83.33% | 13 | 56.52% | 0.090 | 33 | 70.21% |

MMSE = Mini-Mental State Examination; SD = standard deviation.

### MRI Data Acquisition and Preprocessing

Structural MRI scans were obtained using the ADNI3 protocol with 3 Tesla scanners and a magnetization-prepared rapid acquisition gradient echo (MPRAGE) sequence (TR = 2300 ms; TE = minimum full echo; TI = 900 ms; FOV = 208 × 240 × 256 mm; voxel size = 1 mm³). Preprocessing was conducted using FreeSurfer version 6.0 (Dale et al., 1999), including skull stripping, intensity normalization, and segmentation of entorhinal cortex and hippocampus (Fischl et al., 2002, 2004)s. Total intracranial volume (TIV), entorhinal cortex and hippocampus segmentations were extracted from the aparc+aseg.nii files by FreeSurfer.

### Entorhinal-tentorial incisura Channel Segmentation

To quantify anatomical proximity between the EC and the TI, we developed a semi-automated segmentation pipeline combining FreeSurfer-based anatomical priors with seed-based expansion and manual refinement. EC binary masks were first generated using FreeSurfer’s cortical parcellation, and served as anatomical references for identifying the channel space between the EC and TI. The segmentation workflow comprised four stages: medial cluster identification, low-intensity voxel thresholding, seed-based voxel expansion, and guided manual correction. This approach was designed to reduce the influence of noise and segmentation artifacts while preserving anatomical validity across participants.

To initialize the segmentation, we first isolated the two largest contiguous clusters from each hemisphere of the EC mask. These clusters provided stable EC masks by minimizing contributions from fragmented or spurious voxels, especially along the mediolateral boundary of EC. To define the medial portion of the EC in each coronal slice, we identified the most inferior voxel of the EC mask within that slice. A vertical line along the superior-inferior axis was then drawn through this point. EC voxels located medial to this vertical line were selected to form the medial EC mask. The medial EC mask served as a reference to find the starting seed for space searching within the channel between the EC and the TI.

Low-intensity voxels corresponding to the air-filled channel were identified using an adaptive thresholding strategy. Specifically, the mode intensity of EC voxels was computed, and a threshold was defined at 70% of this value to suppress signal contributions from surrounding gray matter and bone. Voxels within a 2-voxel medial range of the medial EC mask and falling below this threshold were designated as seeds. An 8-connected component expansion was then applied to every coronal slice. Each seed voxel was iteratively expanded to its 8-connected neighbors if they satisfied both intensity and spatial constraints. This process produced a preliminary delineation of the channel between the EC and the TI.

Following automatic segmentation, masks were manually refined to ensure anatomical accuracy. Refinement was guided by native-space T1-weighted MRI. Skull stripped T1-weighted MRI can also serve the purpose. Boundaries were adjusted using three constraints. Firstly, channel masks were restricted within the outer boundary of the skull, with air-filled pockets in the skull excluded. Secondly, masks were constrained in-between the lateral boundary of the skull and the medial boundary of the EC. Lastly, voxels within the rhinal sulcus (also known as the olfactory sulcus) were excluded to avoid false-positive labeling. The resulting segmentations provided subject-specific representations of the EC–TI channel suitable for morphometric analysis.

### Neuroanatomical Contact Coefficient Calculation and Anatomical Risk Classification

Individual variation in the EC-TI channel spatial adjacency was quantified using a neuroanatomical contact coefficient (NCC) derived from segmented channel masks and absolute entorhinal volume. For each participant, left-to-right distances between the EC and TI were computed on each coronal slice where both structures were present. These distances were then summed across the full rostrocaudal extent to generate an absolute channel space NCC.

To adjust for differences in overall brain size, the absolute NCC was normalized by the EC volume raised to the 2/3 power, scaling the metric in surface area units. This normalization preserved morphometric interpretability while controlling for individual variability. For finer classification, the resulting proximity values were scaled by a factor of 100. The final output, termed the neuroanatomical contact coefficient, served as a continuous measure of EC–TI adjacency.

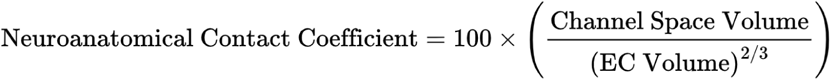

Based on NCC, participants were stratified into anatomical risk groups. Hemispheric indices above 500 were labeled hemispheric low-risk, while values at or below 500 were classified as hemispheric high-risk. Individuals were considered high-risk overall only if both hemispheres met the high-risk criterion, while all others were categorized as low-risk. This binary classification was used in subsequent statistical models to test whether EC–TI proximity modulated the effects of tau pathology, regional atrophy, or conversion to Alzheimer’s disease.

### Relative Entorhinal Cortex and Hippocampus Volume Calculation

To account for individual variance, relative volumes were calculated by dividing each by the total intracranial volume (TIV), resulting in the entorhinal cortex-TIV (EC-TIV) and the hippocampus-TIV (Hippo-TIV) ratios. Regional volumes were extracted from masks generated by FreeSurfer 7.3, and TIV was derived as the sum of all volumes within the aparc+aseg.nii file. Hemispheric relative volumes were scaled by a factor of 2 x 10^5^, while the total relative volumes were scaled by a factor of 10^5^ for better distinction.

### Statistical Analysis

Statistical analyses were conducted in Python 3.0 using libraries including NumPy, Pandas, and Matplotlib. Two-sample t-tests were applied to normally distributed continuous variables (age, MMSE score, EC-TIV Ratio, and tau PET SUVr). Chi-square tests were used for comparison in binary variables (sex, PET amyloid status, conversion, and EC anatomical risk). Statistical significance was set at *p* < 0.05. Pearson correlation was used to examine relationships among continuous variables, including relative entorhinal cortex and hippocampus volumes, tau PET SUVr, and the neuroanatomical contact coefficient. Multivariate logistic regressions were conducted within risk groups, with variable combinations to identify predictors of MCI-to-AD conversion. Significance was determined at *p* < 0.05 for individual predictors and at *p* < 0.05 for the model’s likelihood-ratio test (LLR p-value).

## RESULTS

### Baseline Clinical and Demographic Characteristics of Conversion Groups

In summary of the baseline demographic and clinical characteristics of participants, we tested the data grouped by two criteria respectively: 1) whether or not they converted from MCI to AD within 36 months of initial diagnosis, 2) the EC anatomical risk classification carried out in our study.

Among the 47 participants, 24 (51%) progressed to AD within 36 months, while 23 did not (Table 1). Baseline age (74.13 vs. 74.30 years; p = 0.94), sex distribution (50.0% vs. 58.3% female; p = 0.65), and MMSE scores did not differ between converters and non-converters.

### Baseline Clinical and Demographic Characteristics of the EC Anatomical Risk Groups

To further characterize our EC-based anatomical risk classification, we compared demographic features across high and low-risk groups defined by spatial proximity indices developed for this study. Neuroanatomical contact coefficient is calculated in each hemisphere for every subject to quantify anatomical variation in the spatial relationship between the EC and the TI. This metric was derived by summing the mediolateral distance between EC and TI measured in coronal slices (specifically along the horizontal axis of the coronal scan), then normalizing by EC volume to the ⅔ power to account for differences in cortical surface area. The normalized NCC was scaled by 100 and used to assign lateral anatomical risk. Participants with indices below 500 in both hemispheres were classified as high-risk; otherwise, they were assigned to the low-risk group. Normally, with higher risk, the participant is more likely to have EC close to or abutting TI (Figure 2).

**Figure 2.**
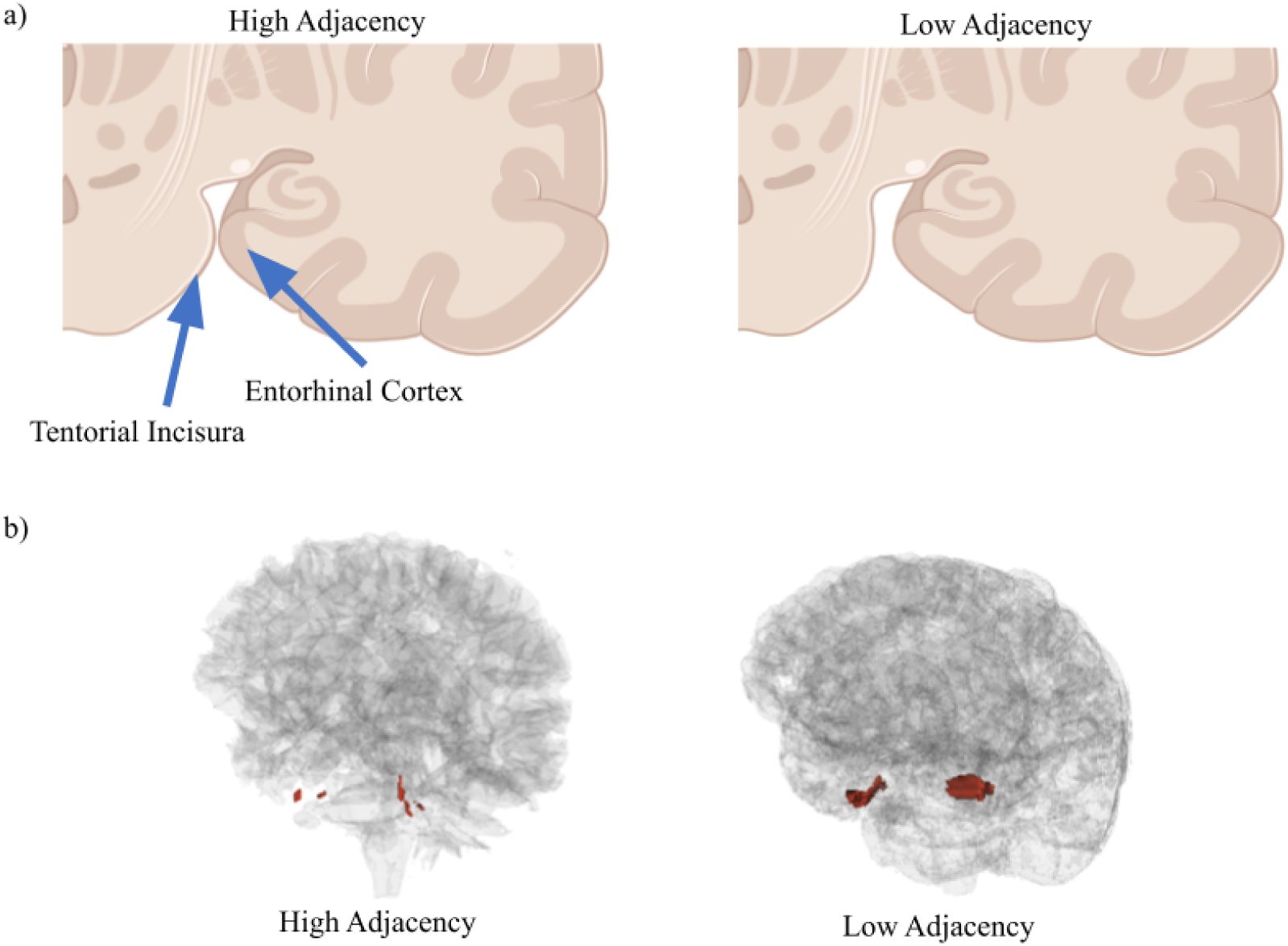
Anatomical classification based on spatial proximity between the entorhinal cortex and tentorial incisura. (a) Conceptual illustration of high-risk (high anatomical adjacency; space proximity index <500) and low-risk (low adjacency; index >500) configurations. (b) Visualization of the channel space between the entorhinal cortex (EC) and tentorial incisura (TI) overlaid on a glass brain. Red regions denote the segmented channel space.

As expected, the left and right proximity indices were significantly (Table 2) higher in the high-risk group (p = 0.0002 and 0.0001, respectively), confirming consistency in the classification approach. Despite anatomical differences, the high- and low-risk groups did not differ in baseline age (73.45 vs. 75.12 years, p = 0.48), sex (58.3% vs. 30.4% female, p = 0.89), MMSE score (26.83 vs. 26.35, p = 0.47), or amyloid positivity (66.7% vs. 73.9%). More importantly, the proportion of participants who converted to AD also showed no difference between groups (54.2% vs. 47.8%, p = 0.89). In conclusion, while proximity-based risk groups reflect anatomical variation, they do not show distinction with clinical features or conversion outcomes.

**Table 2.** Demographic Data, Clinical Assessment, Left and Right Space Proximity Index and Conversion Rates of Participants (n=47)

|  | High Space Risk (n=24) |  | Low Space Risk (n=23) |  | P-value | Total |  |
| --- | --- | --- | --- | --- | --- | --- | --- |
|  | n/Mean | %/SD | n/Mean | %/SD |  | n/Mean | %/SD |
| Baseline Age, years | 73.45 | 8.04 | 75.12 | 7.16 | 0.483 | 74.21 | 7.58 |
| Female, n | 14 | 58.33% | 7 | 30.43% | 0.886 | 21 | 44.68% |
| MMSE | 26.83 | 1.83 | 26.35 | 2.66 | 0.468 | 26.60 | 2.26 |
| Amyloid Positive, n | 16 | 66.67% | 17 | 73.91% | 0.823 | 33 | 70.21% |
| Left Space Proximity Index | 44.90 | 30.04 | 181.30 | 164.20 | <0.0005 | 111.65 | 134.52 |
| Right Space Proximity Index | 32.98 | 24.42 | 119.67 | 98.56 | <0.0005 | 75.40 | 82.84 |
| Conversion, n | 13 | 54.17% | 11 | 47.83% | 0.886 | 24 | 51.06% |

MMSE = Mini-Mental State Examination; SD = standard deviation.

### Space Risk Affects Prediction of Conversion to AD by tau PET SUVr

### Regional Brain Volume and Tauopathy Differences Across Space Risk Groups

Focusing on the EC anatomical stratification, we further evaluate its biological relevance, specifically, we compared the relative EC and hippocampus (HC) volumes, regional tau PET SUVr, between high- and low-risk groups defined by bi-lateral spatial proximity indices. Among lateral and overall EC and HC volumes, only the left hippocampus showed a significant difference (Table 3). Subjects in the high-risk group had larger left HC volumes compared to those in low-risk group (694.05 ± 98.40 vs. 633.93 ± 104.93; *p* = 0.049). Similarly, tau PET SUVr in both the EC and HC on the left and right side did not differ significantly between risk groups.

**Table 3.** Imaging Metrics Comparison between High Channel Risk Group and Low Channel Risk Group.

|  | High Channel Space Risk Group (n=24) |  |  |  | Low Channel Space Risk Group (n=23) |  |  |  | P-value |
| --- | --- | --- | --- | --- | --- | --- | --- | --- | --- |
|  | Mean | SD | L95%CI | U95%CI | Mean | SD | L95%CI | U95%CI |  |
| **EC size** |  |  |  |  |  |  |  |  |  |
| EC/TIV Ratio (x 10<sup>5</sup>) | 294.36 | 50.54 | 273.58 | 315.14 | 288.20 | 61.34 | 262.44 | 313.96 | 0.709 |
| LEC/TIV Ratio (x 2 x 10<sup>5</sup>) | 306.68 | 62.89 | 280.83 | 332.54 | 290.24 | 62.32 | 264.06 | 316.41 | 0.373 |
| REC/TIV Ratio (x 2 x 10<sup>5</sup>) | 282.03 | 50.81 | 261.14 | 302.92 | 286.17 | 78.87 | 253.04 | 319.29 | 0.831 |
| **HC size** |  |  |  |  |  |  |  |  |  |
| HC/TIV Ratio (x 10<sup>5</sup>) | 687.18 | 88.60 | 650.75 | 723.60 | 648.04 | 69.15 | 619.00 | 677.08 | 0.099 |
| LHC/TIV Ratio (x 2 x 10<sup>5</sup>) | 694.05 | 98.40 | 653.60 | 734.51 | 633.93 | 104.93 | 589.87 | 678.00 | 0.049 |
| RHC/TIV Ratio (x 2 x 10<sup>5</sup>) | 682.35 | 111.38 | 636.56 | 728.14 | 672.24 | 103.26 | 628.87 | 715.61 | 0.749 |
| **Tau SUVr** |  |  |  |  |  |  |  |  |  |
| tau PET SUVr in EC | 1.440 | 0.301 | 1.316 | 1.563 | 1.474 | 0.302 | 1.347 | 1.601 | 0.700 |
| tau PET SUVr in HC | 1.348 | 0.221 | 1.258 | 1.439 | 1.393 | 0.214 | 1.303 | 1.483 | 0.486 |

L95%CI = lower 95% confidence interval; U95%CI = upper 95% confidence interval; SD = standard deviation; EC = entorhinal cortex; LEC = left entorhinal cortex; REC = right entorhinal cortex; HC = hippocampus; LHC = left hippocampus; RHC = right hippocampus; TIV = total intracranial volume; PET = Positron emission tomography; SUVr = standardized uptake value ratio.

We examined whether regional brain volume was associated with tauopathy in the EC and HC. No significant correlation was found between relative EC volume (entorhinal-total intracranial volume ratio, EC/TIV Ratio) and EC tau SUVr (Figure 3) in the full cohort (*r* = –0.18, *p* = 0.215), within the high-(*r* = –0.19, *p* = 0.379) or low-risk groups (*r* = –0.18, *p* = 0.417). In contrast, a weak but significant negative correlation emerged between relative hippocampus volume (hippocampus-total intracranial volume ratio, HC/TIV Ratio) and hippocampal tau SUVr (Figure 4) in the full cohort (*r* = –0.33, *p* = 0.023). This relationship was not significant within the high-risk (*r* = –0.34, *p* = 0.103) or low-risk groups (*r* = –0.29, *p* = 0.183).

**Figure 3.**
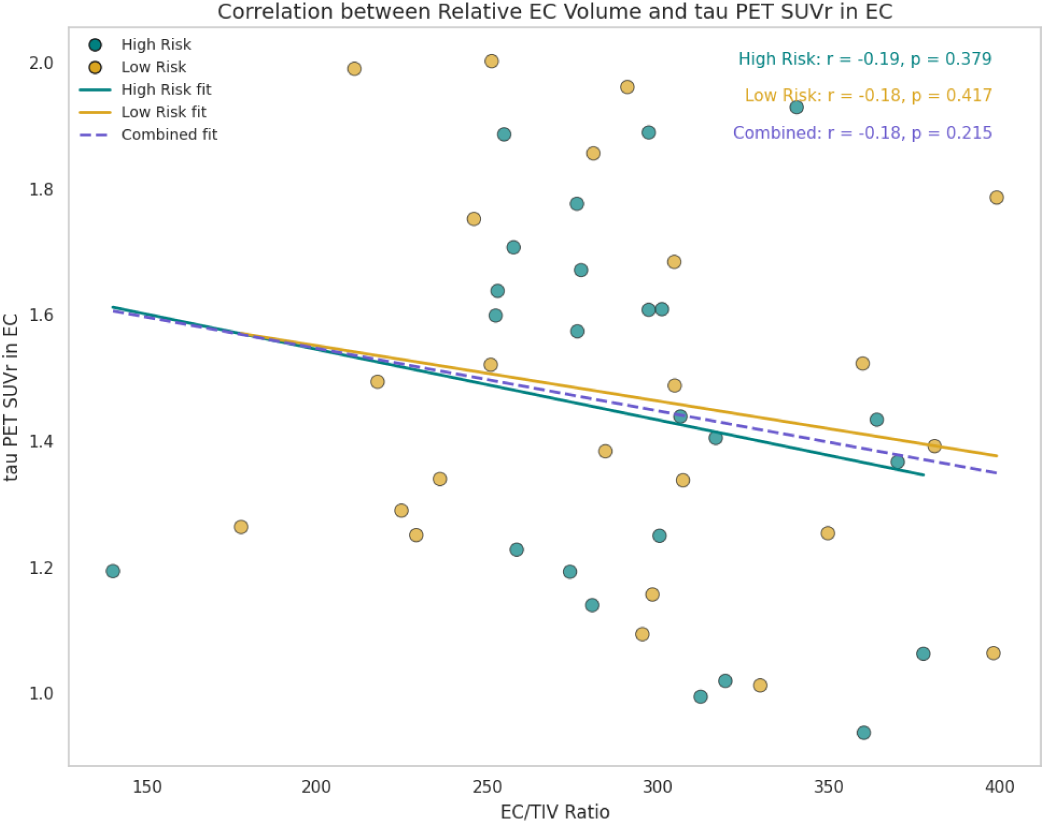
Correlation between Relative EC Volume and tau SUVr in EC

**Figure 4.**
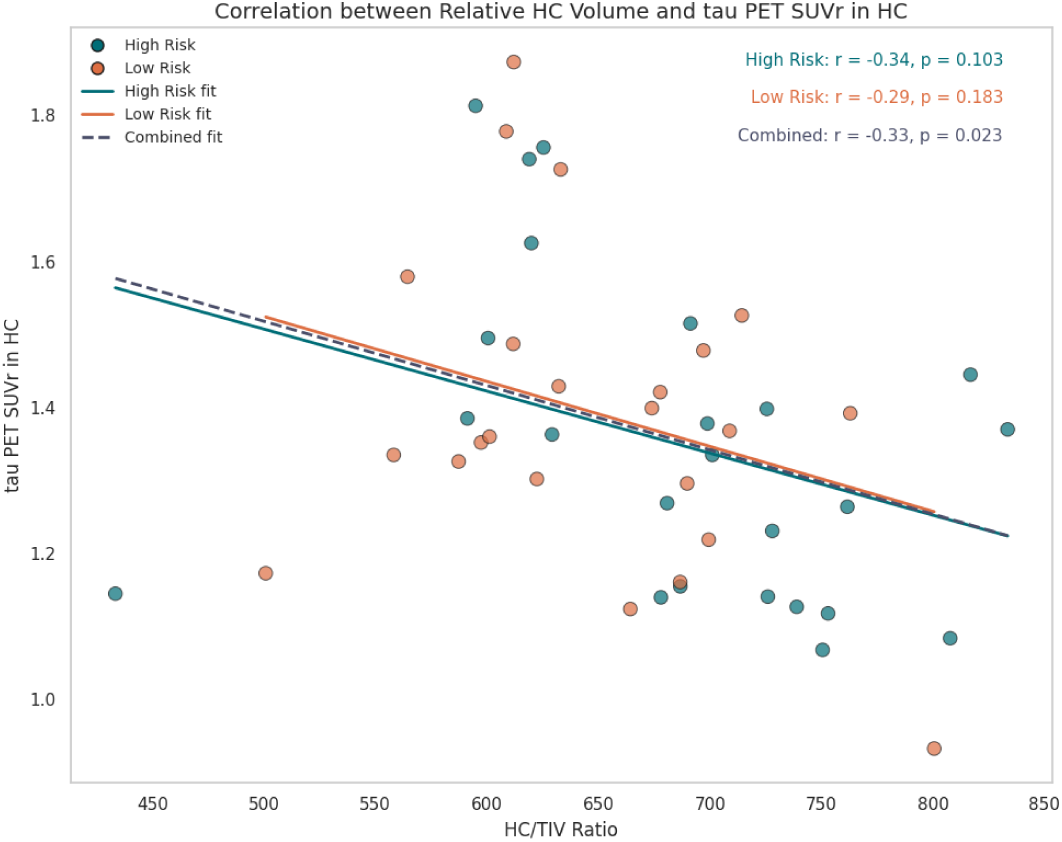
Correlation between Relative HC Volume and tau SUVr in HC

### Multivariate Logistic Regression: Predicting MCI-to-AD Conversion with Anatomical Stratification and Tauopathy

We modeled the within 36 months MCI-to-AD conversion using multivariate logistic regression (Table 4) in the overall cohort, incorporating demographic data, MMSE scores, relative EC and HC volumes, regional tau PET SUVr, and lateral space proximity indices. The model was statistically significant (LLR *p* = 0.012*) with moderate explanatory power (pseudo R² = 0.323). The log-likelihood of the fitted model compared with the null model improved from -32.567 to -22.043.

**Table 4.1.** Logistic Model in Combined Group: Conversion ∼ Age + Sex + MMSE + HC/TIV Ratio + tau SUVr in HC + EC/TIV Ratio + tau SUVr in EC + Left Space Proximity Index + Right Space Proximity Index.

|  | Regression Model |
| --- | --- |
| Pseudo R-square | 0.323 |
| Log Likelihood | -22.043 |
| LL-Null | -32.567 |
| LLR p-value | <0.05 (0.012) |

LL-Null = Log-Likelihood of the Null Model; LLR = Log-Likelihood Ratio.

**Table 4.2.** Estimated Coefficients for the Predictors to Predict Conversion from MCI to AD.

| Variables | Coeff.($\beta$) | Std. Err. | Z-value | P-value |
| --- | --- | --- | --- | --- |
| Const. | -4.33 | 7.73 | -0.56 | 0.576 |
| Age | -0.02 | 0.05 | -0.33 | 0.739 |
| Sex | 1.42 | 1.02 | 1.39 | 0.165 |
| MMSE | -0.02 | 0.18 | -0.13 | 0.894 |
| HC/TIV Ratio | -0.01 | 0.006 | -1.51 | 0.131 |
| tau SUVR in HC | -5.47 | 3.49 | -1.57 | 0.117 |
| EC/TIV Ratio | 0.03 | 0.01 | 2.60 | <0.01 (0.0094) |
| tau SUVR in EC | 7.91 | 2.97 | 2.66 | <0.01 (0.0078) |
| Left Space Proximity Index | -0.02 | 0.01 | -2.20 | <0.05 (0.0279) |
| Right Space Proximity Index | 0.02 | 0.008 | 2.28 | <0.05 (0.0224) |

Coeff.( $\beta$ ) = regression coefficient; Std. Err. = standard error; Const. = Constant; MMSE = Mini-Mental State Examination; HC = hippocampus; EC = entorhinal cortex; TIV = total intracranial volume; PET = Positron emission tomography; SUVR = standardized uptake value ratio.

Age, sex, MMSE scores, relative hippocampal ratio, and HC tau SUVr were not significant predictors (*p* > 0.1 for all). In contrast, entorhinal tau SUVr significantly predicted the MCI-to-AD conversion, emphasizing the link between tauopathy and AD progression. The EC/TIV ratio was also significant (β = 0.03, *p* = 0.0094*), indicating that greater relative EC volume was associated with increased MCI-to-AD conversion risk. It is worth noting that the left and right space proximity indices showed opposite effects in the regression. Left NCC was associated with reduced conversion risk (β = –0.02, *p* = 0.028*), while right proximity was associated with increased risk (β = 0.02, *p* = 0.022*).

To assess whether the EC anatomical risk classification modulates other predictors of conversion, we performed logistic regression respectively in high and low EC anatomical risk groups. The first model excluded hippocampal metrics (Table 5), while the second included them (Table 6).

**Table 5.1.** Logistic Regression Model in High Risk v.s. Low Risk Group: Conversion ∼ Age + Sex + MMSE + EC/TIV Ratio + tau SUVr in EC.

|  | Regression Model in High Channel Space Risk Group | Regression Model in Low Channel Space Risk Group |
| --- | --- | --- |
| Pseudo R-square | 0.464 | 0.139 |
| Log Likelihood | -8.877 | -13.704 |
| LL-Null | -16.552 | -15.921 |
| LLR p-value | <0.01 (0.009) | 0.489 |

LL-Null = Log-Likelihood of the Null Model; LLR = Log-Likelihood Ratio.

**Table 5.2.** Estimated Coefficients for the Predictors to Predict Conversion from MCI to AD.

| Variables | Regression Model in High Channel Space Risk Group |  |  |  | Regression Model in Low Channel Space Risk Group |  |  |  |
| --- | --- | --- | --- | --- | --- | --- | --- | --- |
| | Coeff. ($\beta$) | Std. Err. | Z-value | P-value | Coeff. ($\beta$) | Std. Err. | Z-value | P-value |
| Const. | 0.28 | 0.85 | 0.33 | 0.740 | 0.26 | 0.93 | 0.28 | 0.778 |
| Age | 0.15 | 0.69 | 0.22 | 0.828 | -0.83 | 0.52 | -1.60 | 0.109 |
| Sex | 0.49 | 1.29 | 0.38 | 0.704 | -0.52 | 1.18 | -0.45 | 0.656 |
| MMSE | -1.25 | 1.02 | -1.22 | 0.223 | 0.38 | 0.48 | 0.80 | 0.423 |
| EC/TIV Ratio | 1.85 | 1.37 | 1.35 | 0.177 | 0.47 | 0.52 | 0.91 | 0.364 |
| tau PET SUVR in EC | 2.66 | 1.27 | 2.10 | <0.05<br>(0.036) | 0.29 | 0.53 | 0.55 | 0.586 |

Coeff.( $\beta$ ) = regression coefficient; Std. Err. = standard error; Const. = Constant; MMSE = Mini-Mental State Examination; HC = hippocampus; EC = entorhinal cortex; TIV = total intracranial volume; PET = Positron emission tomography; SUVR = standardized uptake value ratio.

**Table 6.1.** Logistic Regression Model in High Risk v.s. Low Risk Group: Conversion ∼ Age + Sex + MMSE + HC/TIV Ratio + tau SUVr in HC + EC/TIV Ratio + tau SUVr in EC.

|  | Regression Model in High Channel Space Risk Group | Regression Model in Low Channel Space Risk Group |
| --- | --- | --- |
| Pseudo R-square | 0.579 | 0.476 |
| Log Likelihood | -6.963 | -8.351 |
| LL-Null | -16.552 | -15.921 |
| LLR p-value | <0.01 (0.008) | <0.05 (0.03) |

LL-Null = Log-Likelihood of the Null Model; LLR = Log-Likelihood Ratio.

**Table 6.2.**
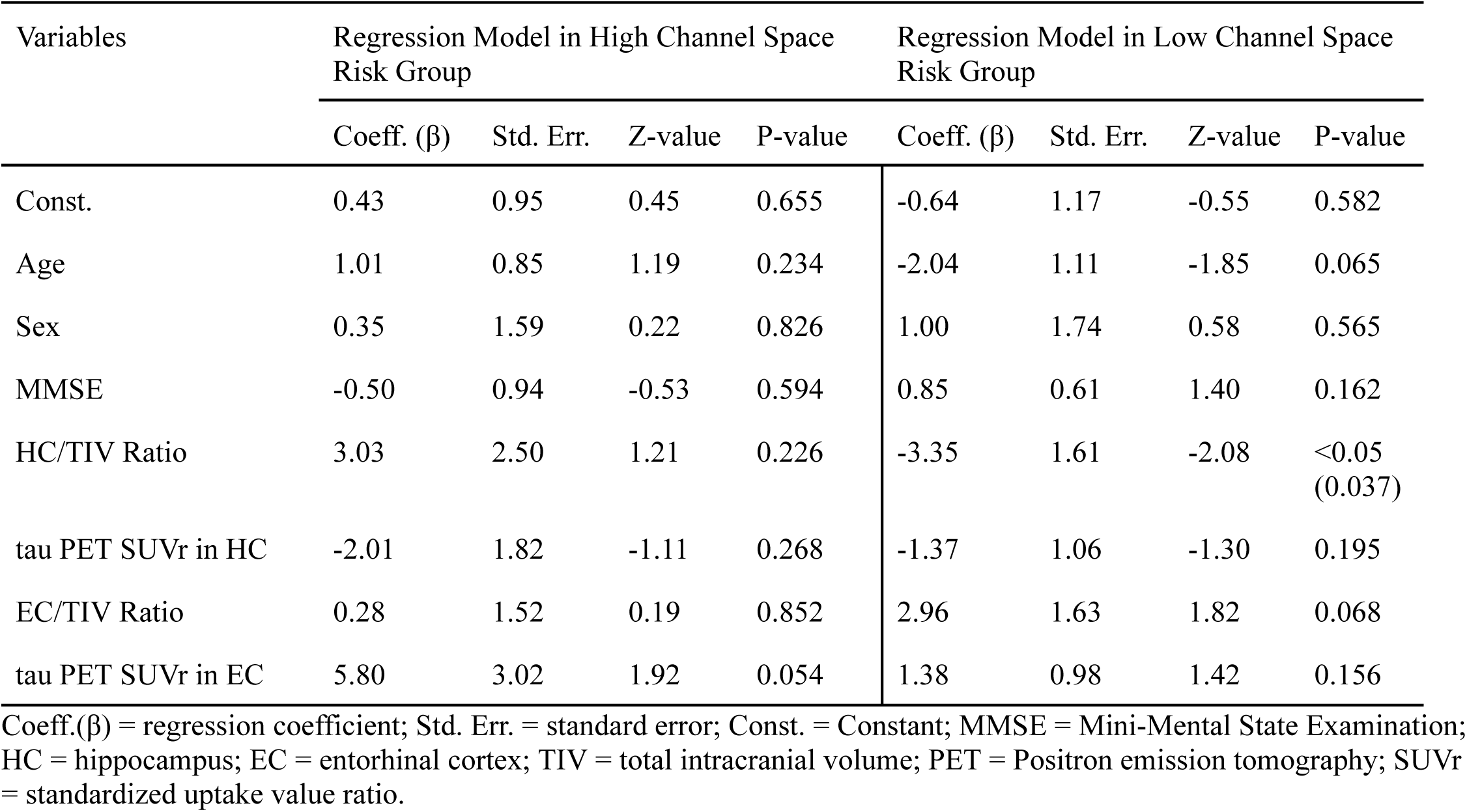
Estimated Coefficients for the Predictors to Predict Conversion from MCI to AD.

| Variables | Regression Model in High Channel Space Risk Group |  |  |  | Regression Model in Low Channel Space Risk Group |  |  |  |
| --- | --- | --- | --- | --- | --- | --- | --- | --- |
| | Coeff. ($\beta$) | Std. Err. | Z-value | P-value | Coeff. ($\beta$) | Std. Err. | Z-value | P-value |
| Const. | 0.43 | 0.95 | 0.45 | 0.655 | -0.64 | 1.17 | -0.55 | 0.582 |
| Age | 1.01 | 0.85 | 1.19 | 0.234 | -2.04 | 1.11 | -1.85 | 0.065 |
| Sex | 0.35 | 1.59 | 0.22 | 0.826 | 1.00 | 1.74 | 0.58 | 0.565 |
| MMSE | -0.50 | 0.94 | -0.53 | 0.594 | 0.85 | 0.61 | 1.40 | 0.162 |
| HC/TIV Ratio | 3.03 | 2.50 | 1.21 | 0.226 | -3.35 | 1.61 | -2.08 | <0.05<br>(0.037) |
| tau PET SUVR in HC | -2.01 | 1.82 | -1.11 | 0.268 | -1.37 | 1.06 | -1.30 | 0.195 |
| EC/TIV Ratio | 0.28 | 1.52 | 0.19 | 0.852 | 2.96 | 1.63 | 1.82 | 0.068 |
| tau PET SUVR in EC | 5.80 | 3.02 | 1.92 | 0.054 | 1.38 | 0.98 | 1.42 | 0.156 |

Coeff.( $\beta$ ) = regression coefficient; Std. Err. = standard error; Const. = Constant; MMSE = Mini-Mental State Examination; HC = hippocampus; EC = entorhinal cortex; TIV = total intracranial volume; PET = Positron emission tomography; SUVR = standardized uptake value ratio.

Strikingly, EC tau burden could only positively predict MCI-to-AD conversion only among participants classified as high anatomical risk. In this group, the first model was significant (pseudo R² = 0.464; LLR *p* = 0.009), with EC tau SUVr emerging as the only significant predictor (β = 2.66, *p* = 0.036). Other variables, including age, sex, MMSE scores, and EC/TIV ratio, were not significant predictors. In contrast, the model was not significant in the low-risk group (pseudo R² = 0.139; LLR *p* = 0.489), and no predictors reached significance.

Adding hippocampal metrics improved prediction in both the high-(pseudo R² = 0.579; LLR *p* = 0.008) and low-risk group (pseudo R² = 0.476; LLR *p* = 0.03). In the high-risk group, EC tau SUVr showed a trend toward significance (β = 5.80, *p* = 0.054), while hippocampal variables remained non-significant. While, notably, in the low-risk group, the HC/TIV ratio was significant (β = –3.35, *p* = 0.037), indicating smaller hippocampal volumes associated with higher risk of conversion.

## DISCUSSIONS

In this study, we are the first to directly evaluate the contribution of chronic mechanical stress to tau burden in the entorhinal cortex and its role in Alzheimer’s disease initiation and progression using multimodal neuroimaging. We hypothesize that the continual contact over decades of the entorhinal cortex against the tentorial incisura may trigger mechanical injury that leads to neurofibrillary degeneration that ultimately may increase the likelihood of acceleration or progression of individuals at high risk of developing Alzheimer’s disease in concert with the intrinsic accumulation of NFT tangles. We examine this through an analysis of a longitudinal Alzheimer’s disease neuroimaging study containing a cohort of individuals with MCI with and without prodromal Alzheimer’s dementia, with both structural MRI and tau-PET imaging. By developing a method to quantify the adjacency of the entorhinal cortex against the structure medial to it, the tentorial incisura, we were able to, for the first time, define the space in this channel, which we refer to as the “uncal fossa”, as well as categorize individuals into high adjacency (‘high-risk’) and low adjacency (‘low-risk’) and examine what other clinical, imaging and biofluidic measures are likely to define prodromal Alzheimer’s disease.

Within this group, our measure of adjacency heuristically split the initial MCI group relatively evenly, reflecting that the MCI population examined demonstrates a sufficient distribution of the variance of the anatomy we plan to test without needing to further balance the data. There is considerable variance in adult brain and skull physiology and anatomy (Matsumura et al., 2022) and the region between these structures has not been previously studied or reported. The baseline imaging used in this study reflects a single timepoint, acquired when a person is already exhibiting some degree of impairment, as well as accumulated a level of pathology either detectable through imaging, biofluidics or some other unbiased measure. The closest analogue to what this study is examining would be head injury (HI) or TBI, however, the degree and chronicity of injury would vary substantially.

TBI has been associated with elevated risk of developing AD (Gottlieb, 2000), with tau distribution following patterns similar to those found in AD (Koerte et al., 2015; Mohamed et al., 2022). In a study examining individuals with MCI/AD impairment with and without a history HI, elevated tau PET was seen in frontal and parietal lobes in those with a HI (Risacher et al., 2021). The pattern of elevated tau signals in this group were less likely to reflect CTE, which generally initiates from perivascular spaces in the neocortex without necessarily targeting the medial temporal structures (Katsumoto et al., 2019). Details of HI, including location and force, are difficult to retrospectively localize without detailed imaging or neuropathology. Furthermore, they reflect a single event leading to suspected injury and vulnerability. We suspect that tauopathic changes due to HI in MCI/AD are separate from those from the chronic subclinical microtrauma that comes from the surface of the EC near the TI in high risk individuals. However, such risks could be cumulative and could act in concert thus exposing the EC to greater focal damage and tau deposition, in addition to the types of mechanisms seen in TBI.

Methods of quantifying and categorizing abnormal tau accumulations in National Football League (NFL) players have suggested phosphorylated tau accumulation related to including monotonic overloads, repetitive contact leading to cyclic fatigue and “creep”, or the patterns and degree of accumulation over time (Horstemeyer et al., 2019). Tau accumulation in these individuals continues non-linearly once they cease playing and enduring potential injury. When comparing isolated TBI to sports players, there is a general decrease in the force and increase in frequency. Extending this paradigm onto our own study, one can appreciate individuals with high EC/TI proximity as having far lower impulse of abutment with orders of magnitude higher frequency. Although the level of impulse necessary to incur injury throughout ones adulthood might be below the threshold for tau accumulation, isolated events or “jolts” that do not cause a loss of consciousness may result in this potential accumulation.

Mechanically, one way to reduce exposure of a contiguous surface to a coarse structure in mechanical design is to place prominences or bumps on that surface (Bayer, 2004). Coincidentally, within the cortex and neocortex there is one area of the brain that has been shown to have uniquely occurring elevations-the entorhinal cortex. These bumps are called “verrucae” and are thought to originate from prominences in the layer II of the EC (Augustinack et al., 2012). Verrucae have been shown to increase in adulthood and disappearance of these verrucae have been shown to correspond to AD pathology. While the EC is particularly vulnerable to AD, one purported evolutionary and mechanistic benefit of raised structures could be the reduction in surface area to the EC to the TI, or alternatively, these prominences being particularly susceptible to subclinical compression may accelerate the cascade of substantially focal tau accumulation.

There is a predicate for singular structures of the brain may be more vulnerable to injury on the basis of their isolated location. For instance, the anterior cingulate is bifurcated bilaterally by the falx cerebri, which is a stiff structure dividing the hemispheres located along the midbrain. Subfalcine herniations, which are the most common form of brain herniation, are the result of areas of the cingulate gyrus extending around the stiff sickle-shaped dura of the falx (Lee et al., 2001). Therefore, the cingulate gyrus is particularly susceptible to TBI and an area implicated in selective attention and cognitive flexibility. Reduced performance and reduced brain volume, though only for one subregion of the anterior cingulate, have been reported in individuals with known TBI compared to controls (Merkley et al., 2013). This suggests that the normal brain structure and vulnerability of a region to a stiff partition or area can have implications of both functional and structural involvement.

This study has important limitations in understanding the time of MRI with respect to disease course, degree of suspected impairment and extrinsic contributing factors of tau burden. It is not clear where specifically along the medio-lateral aspect of the entorhinal cortex tau accumulation occurs relative to its adjacency to the TI. According to Braak staging, the first evidence of NFT pathology in the EC is the lateral aspect (LEC) of the entorhinal cortex, which is likely less proximal to the TI than the medial (Khan et al., 2014). There may be several factors that are responsible for the initiation of EC-related tau accumulation and eventual spread (Vogel et al., 2020). The presence of potential tau accumulation resulting from proximity to the TI within the EC may be additive and not exclusive to the initial deposition of tau in the LEC/transentorhinal cortex (TEC). As such, areas of tau spread, to both the hippocampus and external cortical areas, may be accelerated by the total tau burden within the entire EC circuit (Zhang et al., 2024). Our findings support this because EC/TI adjacency alone did not significantly predict conversion of MCI to AD, however evidence of tau accumulation within the high adjacency group did, indicating that other factors are dominant in establishing likelihood of progression in this population. Furthermore, and somewhat paradoxically, high-adjacency suggests that the EC is likely to be thicker, which would be in contrast to the relatively thinner EC that are seen in those who ultimately go on to develop AD (Pennanen et al., 2004). Additionally, hippocampal volume was not predictive of progression in our MCI group, indicating that obvious structural MRI findings, likely reflective of sufficient neuronal loss, are not sufficient to augur conversion to AD.

Building upon our findings, we propose a “biomechanical cascade hypothesis for Alzheimer’s disease and other tauopathies”. In this model, chronic low-level mechanical stress, particularly at key points of anatomical vulnerability such as the entorhinal cortex, acts as an initiating or accelerating factor in tau pathology. Analogous to principles in orthopedic and cardiovascular pathology, this hypothesis posits that “progressive mechanical failure” where a singular anatomical vulnerability causes redistributed loading and downstream failure of adjacent tissue is a plausible and under-recognized mechanism in neurodegeneration.

In other systems, this pattern is well-documented. For instance, intervertebral disc degeneration often begins with a single compromised disc losing height and hydration, leading to maladaptive stress redistribution that hastens degeneration of neighboring discs, facet joints, and ligaments (Kirnaz et al., 2022). In the rotator cuff, an initial tendon tear results in overloading of the remaining tendons, leading to cascading failures and shoulder instability (Burkhart et al., 1993). ACL tears in the knee destabilize joint biomechanics, increasing the risk for meniscal injury and long-term cartilage damage (Cognetti & Bedi, 2025). Even in cardiovascular failure, damage to a localized region of myocardium, can lead to a compensatory system, eventually resulting in maladaptive remodeling and functional decline (Gersh & Opie, 1984). These human physiological examples share the theme of initial localized failure leading to diffuse and progressive dysfunction through biomechanical propagation.

Applying this paradigm to the brain, the entorhinal cortex, particularly if in high-adjacency contact with the tentorial incisura, may experience chronic microstrain or focal compression that may either trigger or exacerbate tau accumulation. This mechanical nidus may set off a cascade of localized tau seeding, altered neural connectivity, metabolic stress, and eventual tauopathic spread. The variability in the size and shape of the tentorial notch across individuals may further modulate the degree of stress imparted on adjacent structures (Adler & Milhorat, 2002), including not only the entorhinal cortex but potentially the midbrain and brainstem. A narrower or asymmetric tentorial opening may amplify shear or compression, particularly in scenarios of minor trauma, brain shift, or aging-related atrophy.

It is worth asking whether similar biomechanical principles could underlie other sporadic neurodegenerative diseases. For example, in Parkinson’s disease, the substantia nigra lies near a junction vulnerable to rotational shear forces (Arbogast & Margulies, 1998), while in progressive supranuclear palsy (PSP) and corticobasal degeneration (CBD), both tauopathies, the pathology may reflect susceptibility of midline and subcortical structures to biomechanical strain during normal motion or in the context of aging (Kouri et al., 2011). These diseases share certain pathoanatomical features that may reflect a confluence of constitutional anatomical configuration, vulnerability and chronic micro-injury. The possibility that these regions represent multiple foci of stress concentration, each with its own susceptibility to a potential biomechanical cascade, opens a new avenue for understanding the topographic specificity of tauopathies.

While other mechanisms, such as inflammation (Heneka et al., 2015), excitotoxicity (Lewerenz & Maher, 2015), or viral agents (Itzhaki et al., 2016)), are potential contributors to neurodegeneration, their relative absence as primary initiators in other tauopathies strengthens the case for mechanical injury as a plausible root cause or amplifier. Notably, tau accumulation does not commonly occur in purely inflammatory or viral brain diseases, suggesting that a non-immunologic physical mechanism may be uniquely relevant to the pathogenesis of tauopathies. The convergence of anatomical vulnerability, biomechanical strain, and tau pathology warrants deeper exploration in imaging, postmortem, and experimental models.

## CONCLUSIONS

This study suggests that there may be a subclinical of an important area of the brain next to a stiff and coarse structure that may impart additional tau-related risk of developing Alzheimer’s disease. This is due to the confluence of location, EC vulnerability in AD and variance of anatomical brain structure. This study can serve as a way to explore other factors that are only identifiable ante-mortem that may serve as additional biomarkers of resilience and susceptibility in EC tauopathy. As we continue to discover more about cumulative risk factors and disease comorbidity in Alzheimer’s, it is reasonable based on these findings, to hypothesize that EC/TI abutment can lead to a heightened state of tau-specific EC vulnerability that may influence the course of disease. Future studies should examine this in multiple cohorts with molecular imaging and identify the patterns of subtle tau deposition in earlier disease as well as comparing to neuropathology where available. This study, the code and methods necessary to identify this unique structure as well as the relative ubiquity and value of structural MRI, can provide a framework for other diseases where previously unidentified constitutional brain features have yet to be quantified, tested and explored.

## CODE AVAILABILITY

All relevant code and data available upon reasonable request.

## ACKNOWLEDGEMENTS

Data collection and sharing for this project was funded by the Alzheimer’s Disease Neuroimaging Initiative (ADNI) (National Institutes of Health Grant U01 AG024904) and DOD ADNI (Department of Defense award number W81XWH-12-2-0012). ADNI is funded by the National Institute on Aging, the National Institute of Biomedical Imaging and Bioengineering, and through generous contributions from the following: AbbVie, Alzheimer’s Association; Alzheimer’s Drug Discovery Foundation; Araclon Biotech; BioClinica, Inc.; Biogen; Bristol-Myers Squibb Company; CereSpir, Inc.; Cogstate; Eisai Inc.; Elan Pharmaceuticals, Inc.; Eli Lilly and Company; EuroImmun; F. Hoffmann-La Roche Ltd and its affiliated company Genentech, Inc.; Fujirebio; GE Healthcare; IXICO Ltd.; Janssen Alzheimer Immunotherapy Research & Development, LLC.; Johnson & Johnson Pharmaceutical Research & Development LLC.; Lumosity; Lundbeck; Merck & Co., Inc.; Meso Scale Diagnostics, LLC.; NeuroRx Research; Neurotrack Technologies; Novartis Pharmaceuticals Corporation; Pfizer Inc.; Piramal Imaging; Servier; Takeda Pharmaceutical Company; and Transition Therapeutics. The Canadian Institutes of Health Research is providing funds to support ADNI clinical sites in Canada. Private sector contributions are facilitated by the Foundation for the National Institutes of Health (www.fnih.org). The grantee organization is the Northern California Institute for Research and Education, and the study is coordinated by the Alzheimer’s Therapeutic Research Institute at the University of Southern California. ADNI data are disseminated by the Laboratory for Neuro Imaging at theUniversity of Southern California.

